# Ventral tegmental area acetylcholine generates appetitive and aversive incentive motivation

**DOI:** 10.64898/2026.07.09.737546

**Authors:** Selena G. Savatdy, Lingshu Ye, Nomar Serrano Colón, Gabrielle A. Russell, Serena R. Miller, Kurt M. Fraser

**Affiliations:** Department of Psychology, University of Minnesota, Minneapolis, MN 55455

## Abstract

Ventral tegmental area neurons participate in reward learning, motivation, and movement but also have an emerging role in the regulation of aversive behaviors. While these diverse contributions have been attributed to distinct projections of ventral tegmental area neurons, it remains unresolved how these neurons inherit their array of behavioral functions. The ventral tegmental area is rich with acetylcholine receptors and receives dense cholinergic input from the mesopontine tegmentum. While it is known that acetylcholine release in the ventral tegmental area can increase dopamine output we lack an understanding of the scenarios which necessitate ventral tegmental area acetylcholine release. We sought to provide a comprehensive overview of the participation of ventral tegmental area acetylcholine signaling across valence to motivated behavior. Rats were trained on a diverse array of tasks that allowed for the isolation of the contribution of acetylcholine release in the ventral tegmental area to cue- and context-driven behavior. We used intracranial pharmacology to determine the receptor mechanisms that contribute to acetylcholine’s effects. Ultimately, we found that acetylcholine release in the ventral tegmental area is necessary for appetitive and aversive states to become motivationally relevant and spur reward-seeking and threat-avoidance. We propose that this state-dependent contribution allows ventral tegmental area acetylcholine to act as a motivational gate for behavior across valence. This work expands our view of the interaction between neuromodulatory systems in the brain and opens new directions to the understanding of ventral tegmental area neurons in health and disease.

## INTRODUCTION

Ventral tegmental area neurons are essential for movement, learning, and motivated behavior. This is in large part due to the ventral tegmental area being the source of mesocorticolimbic dopamine neurons (Fields et al., 2007; Morales and Margolis, 2017). While historically the ventral tegmental area has been associated with the generation and regulation of reward-related behaviors there is an emerging focus on the contribution of the ventral tegmental area to aversion (Zweifel et al., 2011; Lammel et al., 2014; Warlow et al., 2024). For instance, a specialized subpopulation of ventral tegmental area dopamine neurons is excited by aversive stimuli and in certain downstream projections activation of dopamine neuron terminals promotes aversion (Brischoux et al., 2009; de Jong et al., 2019; Warlow et al., 2024). These divergent functions of ventral tegmental area neurons has been attributed to different embedding of these neurons in sub-circuits that are defined partially by the projection target of any given ventral tegmental area neuron (Juarez and Han, 2016; Morales and Margolis, 2017; de Jong et al., 2022).

The ventral tegmental area receives inputs from a diverse array of sources that release canonical neurotransmitters, neuromodulators, and neuropeptides (Beier et al., 2015; Faget et al., 2016; Morales and Margolis, 2017; Soden et al., 2020, 2023). However, it has remained unclear how these diverse contributions of ventral tegmental area neurons across reward and aversion are inherited from these myriad inputs. Targeted recording has suggested overlapping information encoded by certain direct inputs onto dopamine neurons (Tian et al., 2016), which raises questions about the establishment of the central function of the ventral tegmental area in reward and aversion. Of particular interest is the dense input from mesopontine tegmental neurons that are obligately cholinergic and synapse directly onto both dopamine and non-dopamine neurons of the ventral tegmental area (Garzón et al., 1999; Omelchenko and Sesack, 2005, 2006; Holmstrand et al., 2010; Holmstrand and Sesack, 2011; Xiao et al., 2016). Stimulation of the mesopontine tegmentum can enhance dopamine neuron activity and dopamine release in the striatum which is dependent on cholinergic signaling in the ventral tegmental area (Blaha et al., 1996; Ericson et al., 1998; Mena-Segovia et al., 2008; Xiao et al., 2016). Cholinergic input to the ventral tegmental area has shown to undergo biophysical adaptations after early life stress and stress hormone exposure (Coimbra et al., 2017; Fernandez et al., 2018). While these data suggest that cholinergic input to the ventral tegmental area may be critical to the function of these neurons, evidence for the conditions under which acetylcholine release in the ventral tegmental area is necessary is less well demonstrated.

What is most well-characterized is a necessity for acetylcholine signaling in the ventral tegmental area for of foodor cocaine-paired cues to support conditioned reinforcement (Solecki et al., 2013; Addy et al., 2015; Wickham et al., 2015; Galaj et al., 2017; Nunes et al., 2019, 2023). Conditioned reinforcement is just one psychological component of motivational value that can be attributed to stimuli in the environment and its observation alone prevents inference into the settings by which acetylcholine may contribute to behavior (Cardinal et al., 2002; Berridge, 2004, 2007; Saunders et al., 2015, 2018). Further, there is evidence that acetylcholine in the ventral tegmental area may alter anxiety-like behaviors indicating functions for ventral tegmental acetylcholine across valence (Greba et al., 2000; Small et al., 2016). Given that ventral tegmental area neurons express a diverse array of nicotinic and muscarinic receptors which may differentially contribute to their function we sought to provide a detailed survey of how acetylcholine signaling in the ventral tegmental area drives behavior. We reveal that there is a function for acetylcholine release in the ventral tegmental area that serves a gate for the generation of motivational states across valence. This work opens up new hypotheses and directions for exploring neuromodulatory interactions in diverse behavioral domains.

## RESULTS

### Ventral tegmental area acetylcholine contributes to the contextual control of reward-seeking

We first sought to understand if ventral tegmental area acetylcholine release contributed to Pavlovian conditioned reward-seeking given the prominent role for neurons in this area in the encoding of reward-prediction errors (Keiflin and Janak, 2015; Watabe-Uchida et al., 2017). We trained rats to discriminate between an auditory cue that was paired with the delivery of sucrose reward at its offset, a CS+, from a separate auditory cue that was never associated with reward, a CS- (**Fig. 1A**; **Fig. S1A**). Once rats had acquired discriminated responding we infused either saline, the nicotinic antagonist mecamylamine, or the muscarinic antagonist scopolamine into the ventral tegmental area. We observed that nicotinic antagonism was without effect on responding to CS+ or CS- (Fig. 1B; interaction F_(1, 14)_=2.688, p=0.1247) and did not affect overall activity as assessed with intertrial port entries (**Fig. 1C**; t_13_=1.085, p=0.2963). However, following muscarinic antagonism in the ventral tegmental area rats exhibited increased responding to the CS- (**Fig. 1D**; interaction F_(1, 13)_=5.073, p=0.0422; post hoc p=0.0069), with no effect on responding to the rewarded CS+. In addition, muscarinic antagonism greatly increased the overall number of port entries made throughout the session (**Fig. 1E**; t_13_=2.657, p=0.0198). This pattern of results following scopolamine infusion was suggestive of a loss of contextual control – that is that rats were disorganized in their behavior and returned to inappropriate conditioned responding to a cue for which they had learned would never predict reward (Millan et al., 2015; Fraser and Janak, 2023).

**Figure 1.**
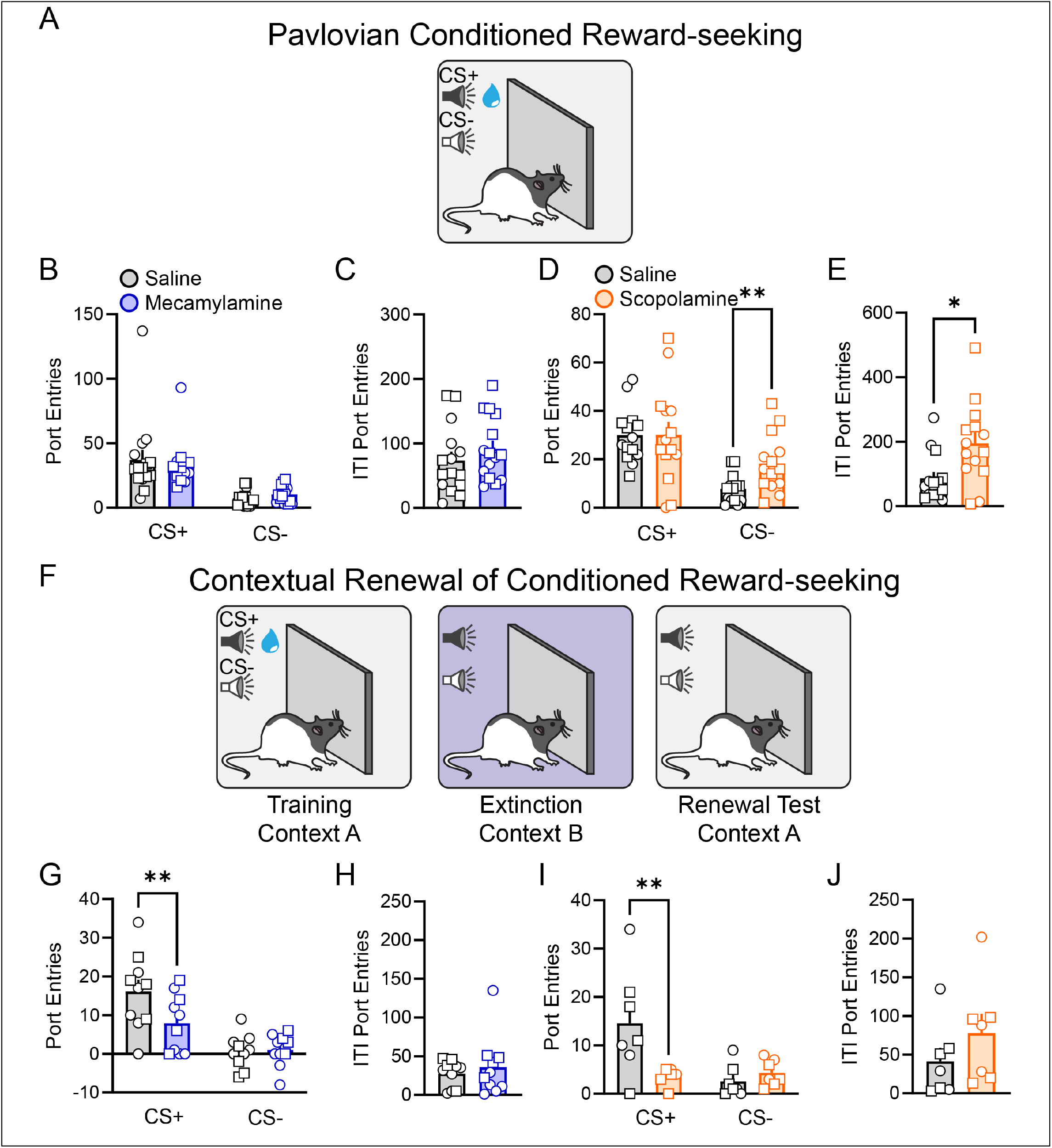
Ventral tegmental area acetylcholine is necessary for reward-paired contexts to energize reward-seeking. A) Rats were trained to associate a 10 s auditory cue with the delivery of 10% sucrose at its offset (CS+) and a separate auditory cue with no reward (CS-). At test rats received infusions of saline, mecamylamine, or scopolamine into the ventral tegmental area. B) Ventral tegmental area nicotinic antagonism did not affect Pavlovian reward-seeking. C) Nicotinic antagonism in the ventral tegmental area did not affect overall activity as assessed by intertrial port entries. D) Muscarinic antagonism within the ventral tegmental area selectively increased inappropriate port entries during the CS-. E) Muscarinic antagonism also significantly increased overall port entry behaviors as evidenced by an increase in intertrial port entries. F) Rats were then moved to a unique context, Context B, and underwent extinction. At test, rats received infusions of saline, mecamylamine, or scopolamine into the ventral tegmental area and then returned to the initial training context, Context A, and presented with the CS+ and CS-but no reward. G) Ventral tegmental area nicotinic antagonism with mecamylamine significantly reduced the renewal of reward-seeking by the reward-paired context. H) Mecamylamine had no influence on port checking behavior during the intertrial interval. I) Ventral tegmental area muscarinic antagonism with scopolamine significantly reduced the renewal of reward-seeking by the reward-paired context and prevented rats from discriminating between the CS+ and CS-. J) Scopolamine did not affect overall port entry behavior as assessed by intertrial port entries. Data presented represent mean + SEM. Individual subjects are overlaid. Circles indicate data from male rats, squares indicate data from female rats. *p<0.05, **p<0.01.

To better isolate if acetylcholine in the ventral tegmental area was contributing to the contextual control of reward-seeking we turned to tests of contextual renewal (Valyear et al., 2020, 2023). We changed the context that the rats were in by altering the smell, flooring, and lighting of the chamber that they were trained in and put the rats through extinction where both the CS+ and CS- were presented but no reward was delivered (**Fig. S1A**). At test, we returned the rats to their original training context after infusions of saline, mecamylamine, or scopolamine into the ventral tegmental area to assay how the motivational value of training context could renew responding to the previously rewarded CS+ (**Fig. 1F**). Critically, at this test no reward is delivered so any responding to the CS+ is due to the motivational renewal of value to just the cue (**Fig. S1B**). Rats in each group demonstrated renewal of reward-seeking at test with higher CS+ than CS-responding following their respective saline infusions (mecamylamine group p=0.0001; scopolamine group p=0.0017). We found that antagonism of either nicotinic (**Fig. 1G**; interaction F_(1,9)_=6.171, p=0.0347) or muscarinic receptors (**Fig. 1I**; interaction F_(1,9)_=6.171, p=0.0347) in the ventral tegmental area disrupted the contextual renewal of reward-seeking. Nicotinic antagonism resulted in a selective blunting of responding to the CS+ (p=0.02). Muscarinic antagonism resulted in a more effective disruption of renewal. Rats no longer responded to the CS+ (p=0.002) and did not discriminate between the CS+ and CS(p=0.7601) after infusions of scopolamine into the ventral tegmental area. Neither mecamylamine (**Fig. 1H**; t_9_=0.6406, p=0.5378) nor scopolamine (**Fig. 1J**; t_6_=2.009, p=0.0913) affected intertrial port entries at test. Thus, acetylcholine release in the ventral tegmental area maintains the motivational properties of rewarding contexts that allows the renewal of cue-driven reward-seeking.

### Ventral tegmental area acetylcholine is necessary for incentive stimuli to enhance reward-seeking actions

Cues paired with reward can come to acquire numerous properties that can motivate behavior. One such motivational property is the ability of a reward-paired cue to energize separately trained reward-seeking actions (Wyvell and Berridge, 2000; Cardinal et al., 2002; Berridge, 2012). This Pavlovian-to-instrumental transfer is reflective of a general motivational process spurred by a state of heightened reward-seeking motivation by reward-paired cues that essentially function as diffuse contexts. Neural activity in the ventral tegmental area, and dopamine neurons therein, is necessary for this motivational effect of reward-paired cues on reward-seeking (Corbit et al., 2007; Halbout et al., 2019). Given that we found that ventral tegmental area acetylcholine diminished the motivational renewal of reward-seeking by contexts, we wondered if in tests of Pavlovian-to-instrumental transfer acetylcholine release in the ventral tegmental area was necessary for the generation of motivational states. We trained rats first to associate a long, 2 minute auditory cue with the random delivery of 20% sucrose (**Fig. S1C**) and then separately trained rats to lever-press for 20% sucrose on a variable interval schedule (**Fig. S1D**). This results in a weak contingency between the Pavlovian cue with the timing and amount of reward received as well as between lever-pressing and reward delivery such that at test under extinction conditions the Pavlovian cue can energize lever-pressing (Malvaez et al., 2026). At test rats received infusions of either saline, mecamylamine, or scopolamine to assess how ventral tegmental area acetylcholine contributed to the generation of states of reward-seeking motivation by reward-cues (**Fig. 2A**). Overall, ventral tegmental area muscarinic antagonism disrupted Pavlovian-to-instrumental transfer by eliminating any elevating effects of cues on lever pressing (**Fig. 2B**; interaction F_(2.904, 52.28)_=4.232, p=0.101) while having little effect on port entry behavior (**Fig. 2E**; treatment F_(1.466, 26.38)_=1.386, p =0.2622)

**Figure 2.**
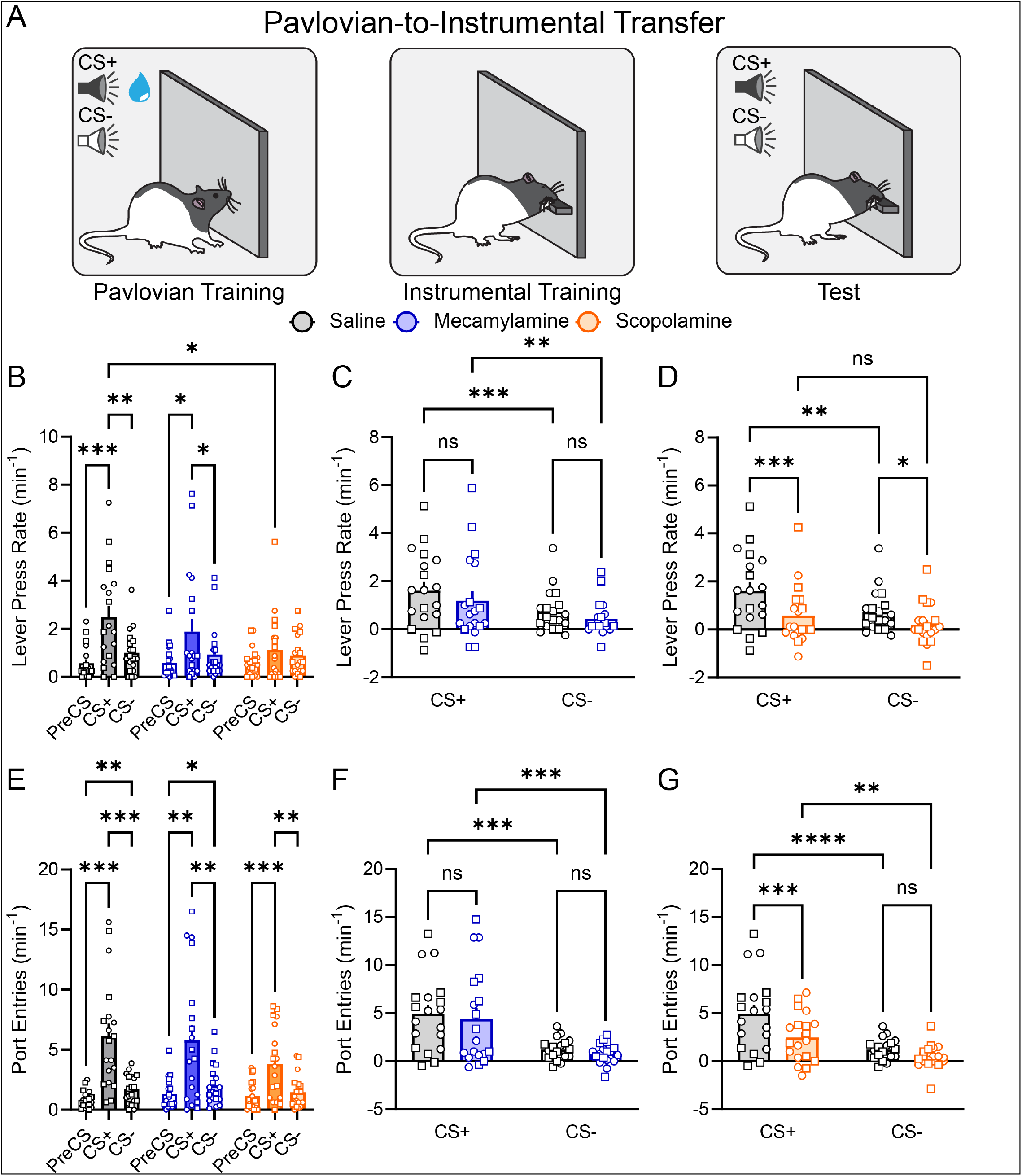
Muscarinic signaling in the ventral tegmental area is necessary for reward-paired cues to generate states of appetitive incentive motivation. A) Rats were trained to separately associate an auditory cue with reward and then separately lever press to earn that same reward. At test, rats were allowed to lever-press in extinction conditions while the reward-paired or control cue were presented. B) Average lever press rates during the pre-cue period, the CS+, and the CS-following ventral tegmental area infusions of saline, mecamylamine, or scopolamine. C) Mecamylamine infused into the ventral tegmental area had no impact on the elevation in lever-pressing elicited by the CS+. D) Scopolamine infused into the ventral tegmental area eliminated the influence of reward-paired cues on energizing reward-seeking actions. E) Average port entry rates during the pre-cue period, the CS+, and the CS-following ventral tegmental area infusions of saline, mecamylamine, or scopolamine. F) Mecamylamine infused into the ventral tegmental area had no impact on the elevation in port-entry behavior elicited by the CS+. G) Scopolamine infused into the ventral tegmental area reduced the influence of reward-paired cues on energizing port entry behaviors relative to baseline at test. Data presented represent mean + SEM. Individual subjects are overlaid. Circles indicate data from male rats, squares indicate data from female rats. *p<0.05, **p<0.01 ***p<0.001, ****p<0.0001.

To better understand these effects we normalized responding for each rat to their behavior in the pre-cue period to assess the elevation in behavior resulting from hearing either the previously rewarded CS+ or the neutral CS-. We found that when rats were tested following scopolamine infusions into the ventral tegmental area, antagonizing muscarinic signaling, they no longer exhibited enhanced lever-seeking actions during the reward-paired cue and also reduced pressing during the control cue (**Fig. 2D**; treatment F_(1,18)_=10.05, p=0.0053; CS+ p=0.0007; CS-p=0.0464). This was in contrast to behavior following nicotinic antagonism where the presentation of the rewarded cue robustly elevated lever-pressing to the same degree as following saline (**Fig. 2C**; cue type F_(1,18)_=7.794, p=0.012; treatment F_(1,18)_=1.619, p=0.2195). In fact, at this test, rats receiving ventral tegmental area scopolamine no longer exhibited any degree of elevation in reward-seeking actions that differentiated behavior between the reward-paired cue and a control cue (saline p=0.0029; scopolamine p=0.1477). Consistent with a loss of cue-generated motivation, scopolamine in the ventral tegmental area also reduced reward cue-evoked port entries (**Fig. 2G**; treatment F_(1,18)_=9.635, p=0.0061; CS+ p=0.0006; CS-p=0.2264). Taken together these data indicate that ventral tegmental area acetylcholine not only maintains motivation elicited by rewarding contexts, but is necessary for reward-paired cues to generate states of appetitive incentive motivation to energize reward-seeking actions.

### Ventral tegmental area muscarinic signaling maintains reward palatability

The prominent role for ventral tegmental area acetylcholine in the motivational value of cues and contexts we observed led us to wonder if acetylcholine release in the ventral tegmental area contributed to primary reinforcement derived from reward (Rada et al., 2000). We first asked if when rats were hungry if ventral tegmental area acetylcholine altered unconditioned motivation to consume a high-value reward - sucrose. We found that ventral tegmental area muscarinic antagonism had a modest effect on sucrose consumption in this brief one-hour test (**Fig. 3C**; t_14_=3.33, p=0.005) but there was no effect of nicotinic antagonism (**Fig. 3B**; t_11_=1.083, p=0.302). To better understand the psychological processes contributing to this change in consumption for a subset of rats we recorded their licking during each test to perform microstructural analysis of reward consumption. We found that the reduction in consumption following ventral tegmental area muscarinic antagonism was due to a significant disruption in the number of licks rats made in a given bout of reward consumption (**Fig. 3E**; t_6_=4.103, p=0.0063), but the overall number of consumption bouts was unaltered (**Fig. 3G**; t_6_=0.3893, p=0.7105). This pattern of changes to rat licking microstructure is indicative of a shift in the perceived palatability of the sucrose (Spector and St John, 1998; Spector et al., 1998; Fraser et al., 2024), but that the rats were still motivated to seek it out and consume the reward. This suggests that ventral tegmental area acetylcholine contributes to reward processing itself but likely is unrelated to generating primary reinforcement.

**Figure 3.**
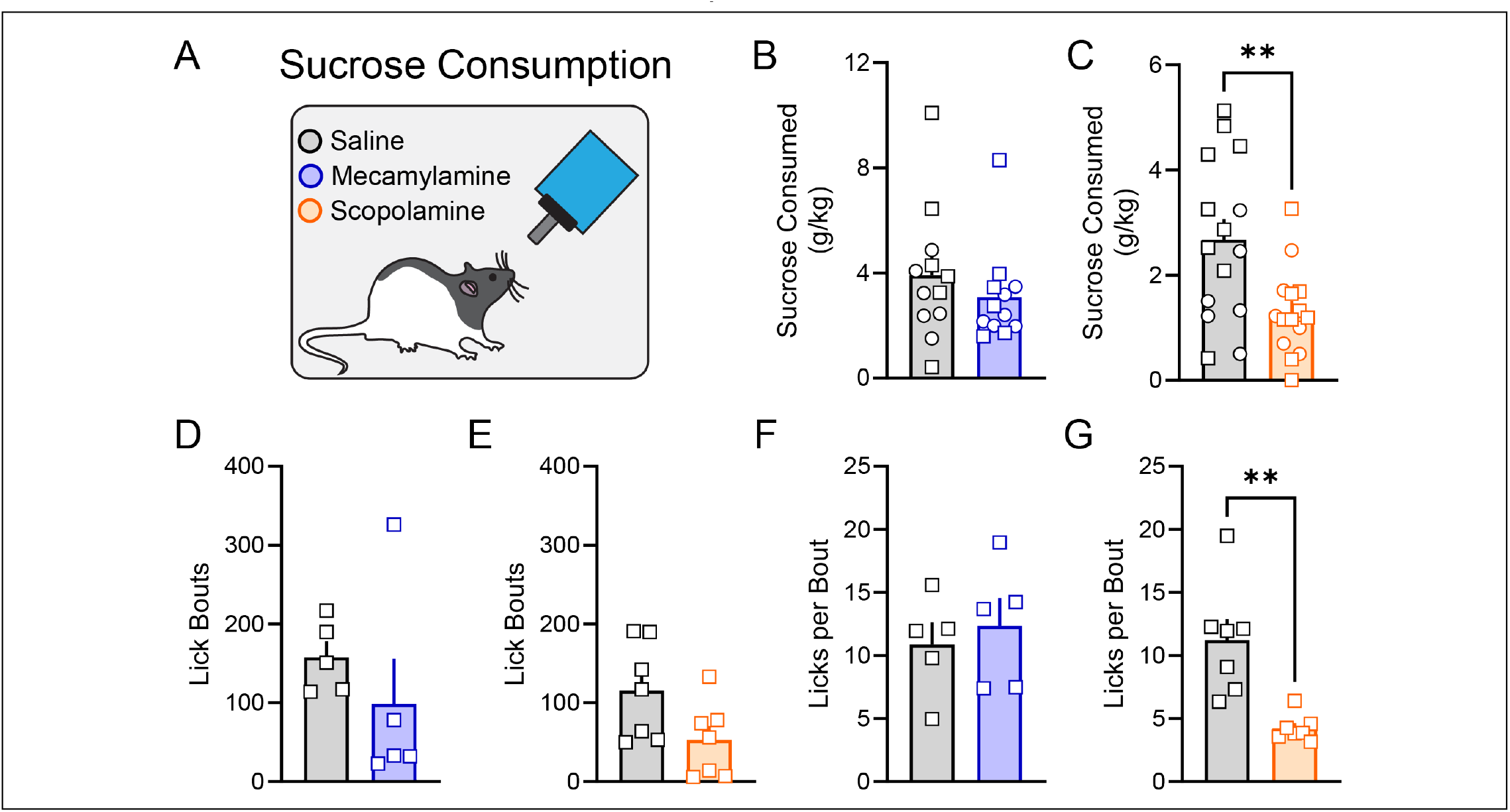
Ventral tegmental area muscarinic signaling maintains the palatability of rewarding outcomes. A) Rats were infused with saline, mecamylamine, or scopolamine into the ventral tegmental area and then allowed to consume 10% sucrose solution for one hour. B) Mecamylamine had no impact on sucrose consumed. C) Ventral tegmental area scopolamine infusions reduced sucrose consumption. D-E) Neither drug infused into the ventral tegmental area altered the number of bouts of sucrose consumption (mecamylamine t_4_=0.8507, p=0.4429). F) Mecamylamine had no impact on the average number of licks in a given bout of sucrose consumption (t_4_=0.9699, p=0.3870) G) Scopolamine significantly reduced the average number of licks in a given bout of sucrose consumption. Data presented represent mean + SEM. Individual subjects are overlaid. Circles indicate data from male rats, squares indicate data from female rats. **p<0.01.

### Ventral tegmental area acetylcholine is dispensable for effortful responding

The reinforcing value of rewards, driven in part by their palatability, can be more sensitively assayed in tasks where the willingness to work for reward is directly assessed. Prior evidence for acetylcholine function in operant responding for reward is mixed (Sharf and Ranaldi, 2006; Sharf et al., 2006; Nunes et al., 2023) We sought to resolve this discrepancy. To do so we trained a group of rats to nosepoke for sucrose reward. We then tested rats in a progressive ratio task where each reward earned increased the subsequent amount of responses required to earn the next reward (**Fig. 4A**). We found that infusions of acetylcholine receptor antagonists into the ventral tegmental area had no effect on motivation as assessed by breakpoint (**Fig. 4C**; F_(1.116, 18.69)_=0.01680, p=0.9184) or total active nosepokes (**Fig. 4D**; F_(1.088, 18.5)_=0.03428, p=0.8736). As with Pavlovian conditioning, we found that ventral tegmental area muscarinic antagonism resulted in disinhibition of inappropriate behaviors as evidenced by higher inactive nose-pokes (**Fig. 4E**; F_(1.785, 30.34)_=7.184, p=0.0037; post hoc p=0.0228) and overall port entries (**Fig. 4F**; F_(1.796, 7.088)_=7.088, p=0.004; post hoc p=0.0393).

**Figure 4.**
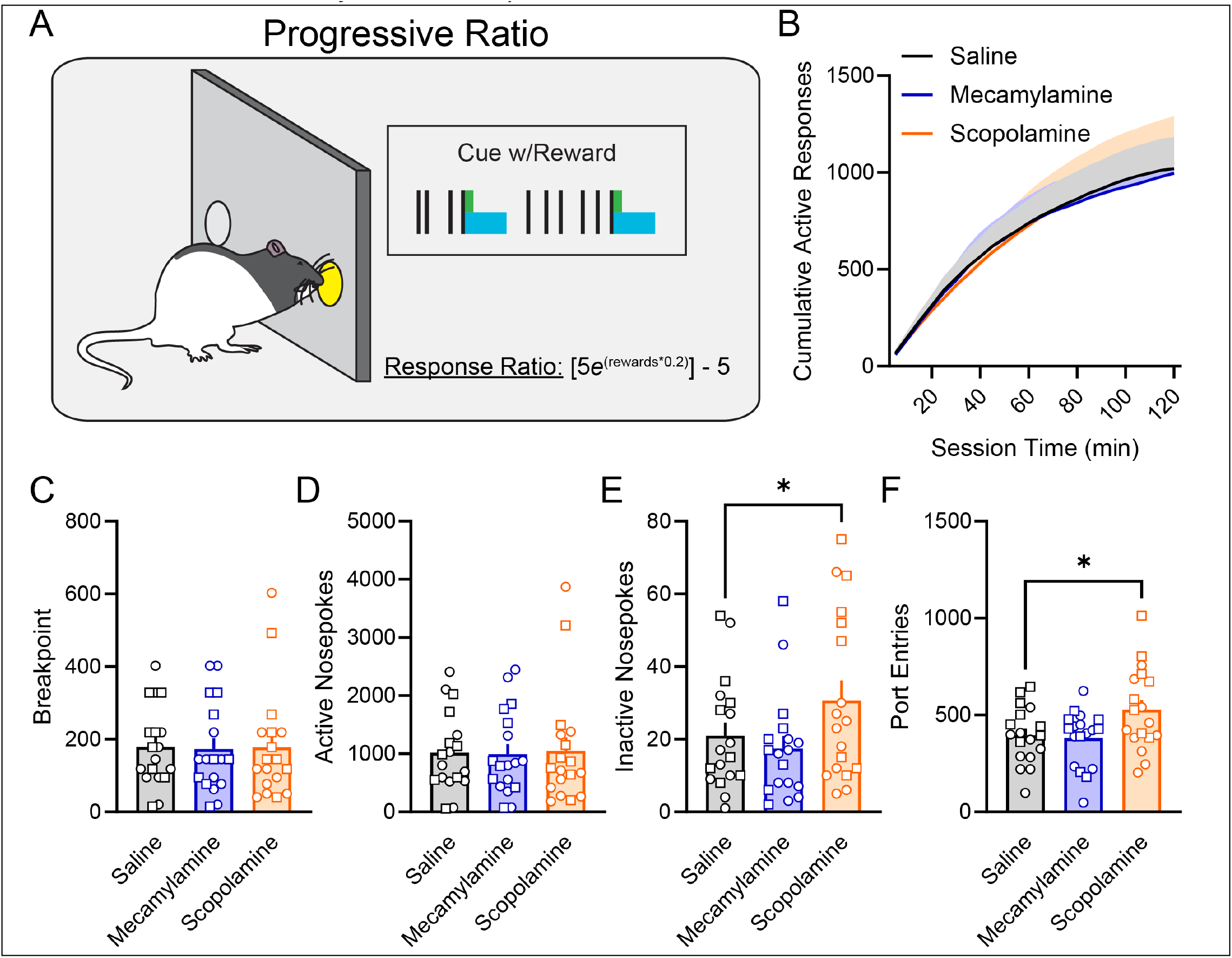
Ventral tegmental area acetylcholine is dispensable for effortful responding for reward. A) Rats were tested on a progressive ratio task where the number of nosepokes required for each reward exponentially increased for each reward earned. B) Cumulative responding at the active nose-poke port was equivalent following ventral tegmental area infusions of saline, mecamylamine, or scopolamine (treatment F_(1.082, 18.39)_=0.004, p=0.9547). C) Manipulation of ventral tegmental area acetylcholine signaling did not affect motivation for sucrose as assessed by breakpoint. D) Active nosepoke responding was unaffected by ventral tegmental area acetylcholine blockade. E) Ventral tegmental area muscarinic antagonism elevated inactive nosepoke responding and F) overall port entries at test. Data presented represent mean + SEM. Individual subjects are overlaid. Circles indicate data from male rats, squares indicate data from female rats. *p<0.05.

These data seem to exclude a role for ventral tegmental area acetylcholine in either the direct reinforcement derived from reward and in the motivation to work for reward itself.

### Acetylcholine in the ventral tegmental area enables contexts to acquire aversive motivational value

Our findings thus far make it apparent that ventral tegmental area acetylcholine contributes to the motivational properties of rewarding contexts and in particular their ability to energize reward-seeking. A challenge with reward conditioning is that the acquisition of learning can take many sessions which impedes the ability to manipulate either reward cue or context learning as it occurs. We wanted to assay the function of ventral tegmental area acetylcholine in the establishment of contexts as motivational generators and turned to threat conditioning to do so. We used Pavlovian cued-threat conditioning where, in a single session, rats acquire conditioned freezing to a 10 s auditory stimulus paired with an aversive 2 s, 1.0 mA foot-shock. This allowed us to antagonize either ventral tegmental area nicotinic receptors or muscarinic receptors as learning occurred to later probe the content of learning in tests for cue memory or context memory. We infused either saline, mecamylamine, or scopolamine into the ventral tegmental area just before threat conditioning so that learning occurred under the effects of the drug. We found that neither drug had an impact in the acquisition of conditioned freezing to the auditory cue during learning (**Fig. 5B,E;** trial F_(3.477, 66.06)_=57.21, p<0.0001; treatment F_(2, 20)_=2.188, p=0.1382). The next day we returned rats, drug-free, to the context in which conditioning had occurred to assay memory for the aversive context in a brief 10 minute test. Surprisingly, we observed that rats who had muscarinic antagonism in the ventral tegmental area during learning had significantly lower freezing at this test compared to rats that had previously received saline or mecamylamine (**Fig. 5C,F;** treatment F_(2,20)_=7.762, p=0.0032; scopolamine p=0.0073). However, when we moved rats to a unique context with lighting, flooring, and smells distinct from the conditioning context and presented the shock-paired cue, all rats exhibited equivalent levels of conditioned freezing (**Fig. 5D,G**; treatment F_(2,20)_=1.023, p=0.3776). As a result, the rats undergoing ventral tegmental area muscarinic antagonism during conditioning had learned the cue-shock association just fine but failed to attribute aversive motivational value to the context where that learning occurred.

**Figure 5.**
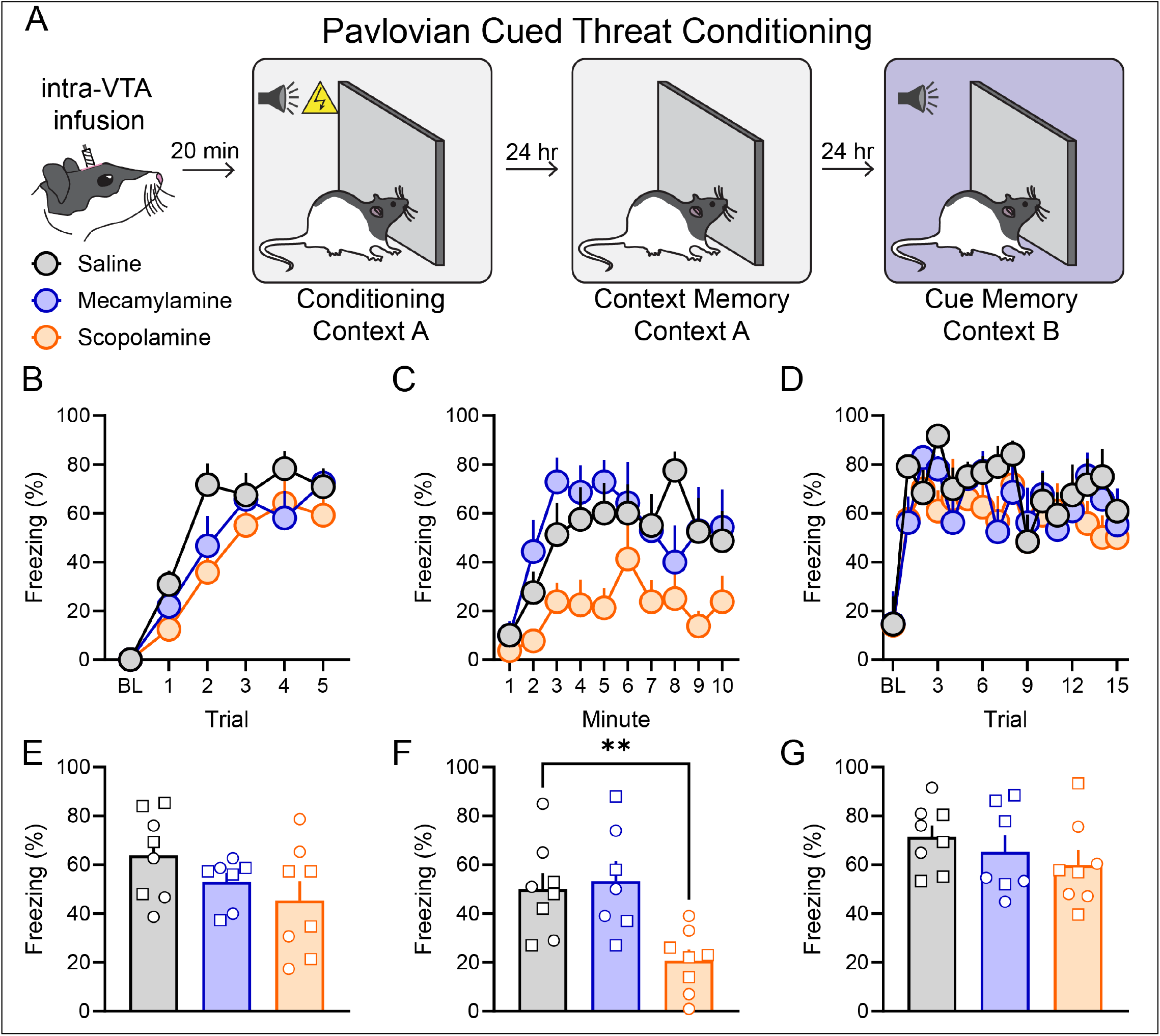
Ventral tegmental area acetylcholine is necessary for threat-paired contexts to acquire aversive motivational value. A) Rats received infusions of saline, mecamylamine, or scopolamine into the ventral tegmental area and then underwent Pavlovian cued threat conditioning where a 10 s auditory cue was paired with footshock. We then assessed rats’ memory for the threat-paired context and threat-paired cue in separate tests while rats were drug-free. B) Ventral tegmental area acetylcholine blockade did not influence the acquisition of cued-threat. C) Antagonism of ventral tegmental area muscarinic receptors during threat learning prevented the threat-associated context from eliciting conditioned freezing. D) Expression of threat to the threat-paired conditioned cue was unaffected by training under ventral tegmental area acetylcholine receptor blockade. E-G) Average freezing behavior during conditioning, context memory test, and the cue memory test. Data presented represent mean + SEM. Individual subjects are overlaid. Circles indicate data from male rats, squares indicate data from female rats. BL=baseline. **p<0.01.

### Ventral tegmental area acetylcholine controls aversive predictions

We next wanted to assess if ventral tegmental area acetylcholine was necessary for the ongoing expression of threat. In our previous experiment one concern is that threat to the cue may be at a maximal level and we would be unable to assess nuances in the expression of threat. To better isolate threat discrimination we used a probabilistic conditioned suppression task which is highly sensitive to detecting changes in behavior driven by an expectation of aversive foot shock (Chu et al., 2024). We trained rats to nosepoke at a high rate for reward on a variable interval schedule while being presented with three distinct auditory stimuli that had a set probability of delivering a 0.5 s, 0.3 mA footshock. One cue always predicted footshock, one cue had a 25% probability of footshock, and the third cue was never associated with footshock. In this task, threat is assessed by the reduction in the rate of reward-seeking elicited by the presentation of each cue (**Fig. S1E-F**). Following training we infused either saline, mecamylamine, or scopolamine and assessed how ventral tegmental area acetylcholine contributed to the ongoing scaling of cue-elicited threat. We found that after ventral tegmental area mecamylamine infusions rats exhibited a significant decrease in overall nose poke rate (**Fig. 6C**; t_21_=2.759, p=0.0118) but there was no effect on expression of threat as assessed by suppression ratios (**Fig. 6B**; F_(1, 21)_=0.3837, p=0.5423; cue discrimination F_(1.76, 36.95)_=39.68, p<0.0001). Ventral tegmental area scopolamine produced a similar decrease in baseline nosepoke rate (**Fig. 6E**; t_21_=5.152, p<0.0001) that was also accompanied by an increase in suppression ratios across all cues (**Fig. 6D**; treatment F_(1, 21)_=4.739, p=0.0410; cue discrimination F_(1.832, 38.46)_=46.49, p<0.0001). Because the suppression ratio is normalized by each rats ongoing nosepoke rate it is relatively resistant to such changes. Instead the nosepoke rate change suggests that following acetylcholine antagonism in the ventral tegmental area there were two distinct effects: 1) inflation of the aversive motivation of the context that reduced reward-seeking actions and 2) muscarinic signaling is essential for producing accurate expectations of threat generated by predictive cues. Together these data implicate ventral tegmental area acetylcholine in balancing reward-seeking and threat-avoidance in mixed valence contexts.

**Figure 6.**
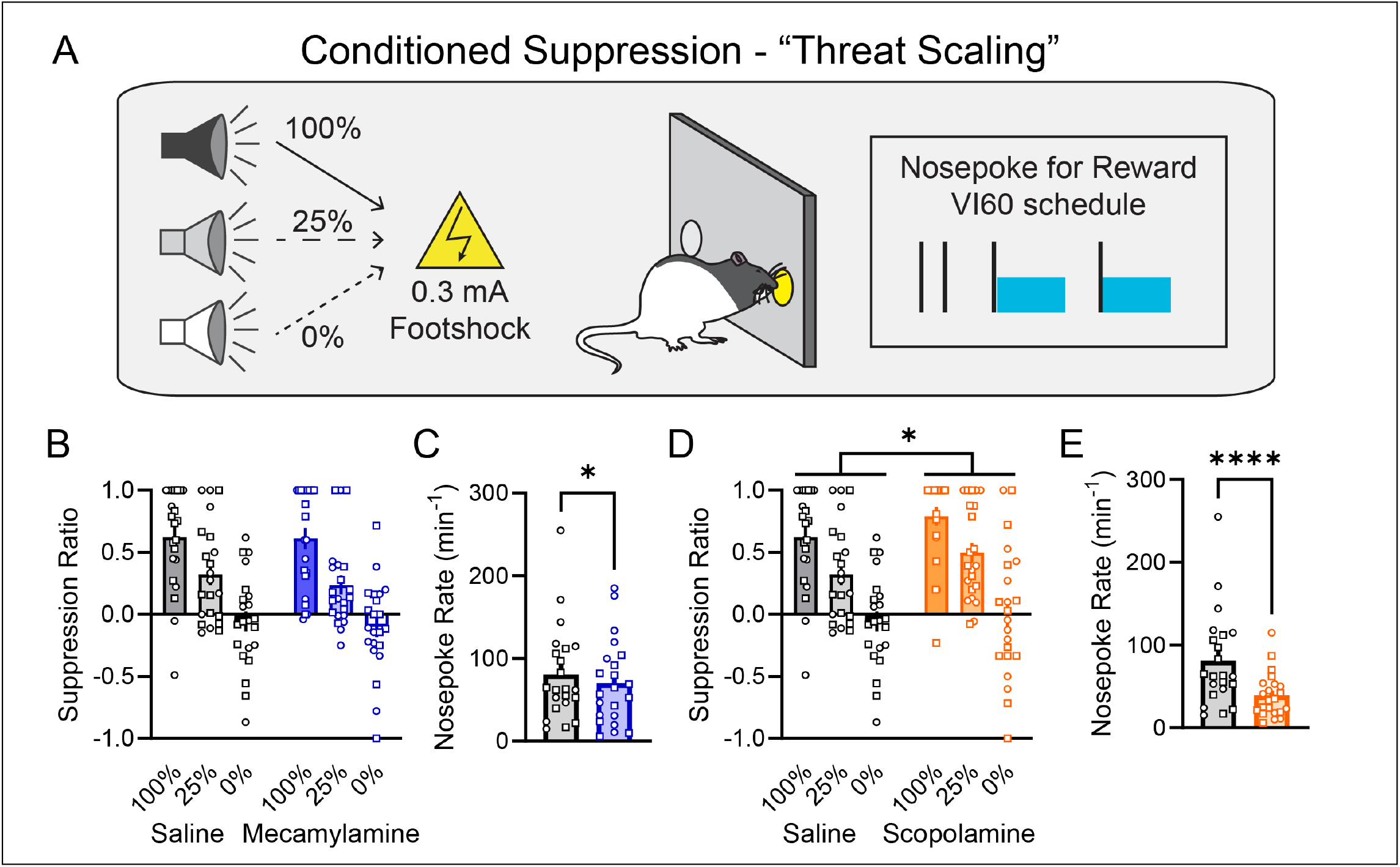
Ventral tegmental area acetylcholine constrains threat expression. A) Rats were trained to distinguish between three auditory cues with defined probabilities of delivering aversive footshock at their offset against a background of nosepoking on a variable interval 60 s schedule for sucrose reward. B) Mecamylamine infused in the ventral tegmental area did not affect threat scaling. C) Mecamylamine in the ventral tegmental area reduced baseline nose poke rates. D) Scopolamine infused in the ventral tegmental area increased threat non-discriminately across all cues. E) Scopolamine also significantly reduced baseline nose poke rate. Data presented represent mean + SEM. Individual subjects are overlaid. Circles indicate data from male rats, squares indicate data from female rats. *p<0.05, ****p<0.0001.

## DISCUSSION

The ventral tegmental area is a critical neural substrate for learning, motivation, and movement. For decades intense focus on the functions of ventral tegmental area neurons has revealed their encoding properties in an array of tasks and their divergent contribution to behavior by their cell-type and projection target. Here we reveal that specialized cholinergic input into the ventral tegmental area is necessary for shaping the behavioral function of this region across reward and aversion. We reveal through a battery of well-controlled tasks that ventral tegmental area acetylcholine release is required for motivational invigoration by rewarding and aversive cues and contexts. This motivational gating by acetylcholine release in the ventral tegmental area across valence reveals novel neuromodulatory interactions of relevance to health and disease.

The ventral tegmental area receives dense cholinergic input exclusively from the mesopontine tegmentum (Holmstrand and Sesack, 2011). These cholinergic inputs lack machinery for co-release of other neurotransmitters and synapse directly onto both dopamine and non-dopamine neurons (Garzón et al., 1999; Omelchenko and Sesack, 2005, 2006; Holmstrand and Sesack, 2011). It has long been established that electrical stimulation of mesopontine nuclei which contain cholinergic neurons elicits robust overflow of dopamine release within the striatum which is blocked by local antagonism of acetylcholine receptors in the ventral tegmental area (Blaha et al., 1996). Further, presentations of cues paired with alcohol elicit striatal dopamine release that is similarly prevented by blockade of ventral tegmental area acetylcholine receptors (Blomqvist et al., 1997; Ericson et al., 1998). It has been proposed that this dense input positions cholinergic nuclei to robustly regulate the dopamine system (Mena-Segovia et al., 2008; Nunes et al., 2024). In agreement, we present here extensive evidence that during states of substantive motivational engagement blockade of ventral tegmental area acetylcholine prevents such motivation. That this function was observed across valence supports a notion by which cholinergic input to the ventral tegmental area functions as a gate for the generation of motivation. Given that we recently detailed a mechanism for such motivational gating supported by dopamine neurons (Fraser et al., 2025), it is tempting to speculate that the source of this function arises from the cholinergic mesopontine tegmentum.

Two nuclei in the mesopontine tegmentum – the pedunculopontine tegmentum and the laterodorsal tegementum – are the source of parallel cholinergic input to the ventral tegmental area.

Activation of either cholinergic input to the ventral tegmental area supports the generation of a conditioned place preference (Xiao et al., 2016; Steidl et al., 2017). Further, long-term inhibition of pedunculopontine cholinergic input to the ventral tegmental area can alter the acquisition of conditioned responding to a reward-paired cue (Yau et al., 2016). However, the underlying psychological processes supported by these parallel cholinergic inputs is not readily inferred from either of these investigations (Saunders et al., 2018; Fraser et al., 2023). Resolving the precise computations and behavioral processes supported by these dual cholinergic mesopontine tegmental nuclei will be critical to advancing our understanding of neuromodulatory interactions.

The incentive motivational properties of cues and contexts – their ability to elicit approach, support conditioned reinforcement, and energize ongoing actions – has long been associated with activity in the mesolimbic dopamine system. Our results suggest that these incentive motivational properties are inherited, in part, by activation of cholinergic inputs in the ventral tegmental area. Further, the aversive motivation elicited by threat-paired cues and contexts appears to similarly require cholinergic activity at muscarinic receptors in the ventral tegmental area. One of the most surprising results we observed was total loss of aversive value attributed to the threat-conditioned context following conditioning under local muscarinic antagonism. A possibility is that the parallel inputs from the laterodorsal and pedunculopontine tegmentum carry distinct motivational signals that separately invigorate appetitive and aversive motivation. In addition to the incentive motivational properties of cues we also observed that muscarinic antagonism reduced the perceived palatability of reward. This is consistent with evidence that reward consumption elevates extracellular concentrations of acetylcholine in the ventral tegmental area (Rada et al., 2000). Despite this, when we asked rats to work on a progressive ratio for sucrose, there was no effect. This dissociation suggests that acetylcholine signaling in the ventral tegmental area may be recruited to help evaluate what is the best available option to potentiate motivation towards the best target. In our studies rats only ever had one reward available so while palatability of that reward may have been affected, its ability to support and goad motivation remained intact.

We consistently observed that muscarinic antagonism within the ventral tegmental area was effective at disrupting appetitive and aversive motivation. In contrast, nicotinic antagonism seemed limited to the motivational influence of reward-paired contexts in energizing responding to reward-paired cues. This difference between nicotinic and muscarinic receptor influencing behavior has also been observed in cocaine-related behavior where nicotinic antagonism seems limited to influence reinstatement for cocaine-paired, but not food-paired, cues (Solecki et al., 2013; Addy et al., 2015; Wickham et al., 2015; Nunes et al., 2019). The ventral tegmental area is one of few regions in the central nervous system that expresses the M5 receptor which makes it a promising target for drug development (Garzón and Pickel, 2013; Razidlo et al., 2022; Nunes et al., 2024). Indeed, we hope to pursue in future work the specific acetylcholine receptors that participate in the generation of appetitive and aversive motivation.

Overall we establish here novel evidence for the involvement of cholinergic systems in regulating and generating incentive motivation across appetitive and aversive states. The cholinergic system is composed of numerous discrete nuclei that likely have distinct psychological and computational functions that are determined by their source and projection target (Baxter et al., 1995; de Cothi et al., 2026; Jang et al., 2026). Targeted manipulations of cholinergic signaling specifically at muscarinic receptors is emerging as a novel treatment for psychiatric illness (Kaul et al., 2024). We reveal here an essential contribution of acetylcholine acting at muscarinic receptors in the ventral tegmental area to the generation of rewarding and fearful motivation. We propose that acetylcholine in the ventral tegmental area specifically acts a gate for motivational generation by cues and contexts. Motivational dysregulation is common to substance use disorders and psychiatric illness so it is our hope that this work provides a launch board for new directions in the field and the development of novel therapeutics.

## MATERIALS & METHODS

### Subjects

Subjects were male and female Long-Evans rats (Envigo; n=66; male n=31; female n=35). Rats were individually housed in a temperature- and humidity-controlled room with a 12:12 light/dark cycle (lights on at 07:00). Rats were allowed to acclimate upon arrival for at least one week before the onset of any experimental procedures. At all times rats had ab libitum access to water and food other than noted below when food restriction (85-90%) occurred during specific behavioral tasks. All experiments occurred during the light cycle. All procedures were approved by the University of Minnesota Institutional Animal Care and Use Committee and in accordance with the Guide for the Care and Use of Laboratory Animals: Eighth Edition, revised in 2011.

For all experiments rats were handled by experimenters for 1-3 days before the onset of experiments. Rats were given 24 hour access to sucrose solutions in the home cage during the handling period to reduce neophobia. Food restriction occurred gradually over 1-3 days. Rats were fed within an hour of the completion of behavioral procedures each day.

Some rats completed more than 1 experiment. Rats undergoing fear conditioning then subsequently had 2 weeks off of behavior before undergoing nosepoke training for conditioned suppression and progressive ratio testing. Rats in the Pavlovian conditioning experiment also underwent reward consumption tests.

### Apparatus

Behavioral training and testing occurred in MedAssociates conditioning chambers (St. Albans, VT) housed in sound- and light-attenuating cabinets and controlled by a computer running MedPC V software. On the center of the wall of the left and right side of each chamber was a fluid receptacle located in a recessed port which allowed for the delivery of liquid rewards. On the right side, the receptacle was flanked on either side by retractable levers. On the left side, the receptacle was flanked by nose poke ports. Rewards were delivered by a 30 mL syringe placed in a motorized pump located outside the cabinet that when activated delivered 0.1 mL of reward per second. Each chamber was equipped with a white noise generator, a solenoid clicker, 2.9 kHz tone generator, and a 4.5 kHz tone generator that all produced their respective auditory stimuli at ∼80 dB when activated. A stainless steel grid floor was connected to a scrambled shock generator that, if activated, could supply shocks directly to the grid floor. Background illumination was supplied by a house light located within the chamber and a light located within the cabinet while background noise was supplied by a fan (70 dB).

For lick recordings during consumption we used custom-designed rat cages that allowed for the presentation of two ball-bearing sipper bottles that hung outside of the cage and were connected to a capacitive sensor (Arduino). Consumption was assessed by weighing bottles and the simultaneous detection of licking via the capacitive sensor. Licks were detected in real-time and logged onto a nearby PC running custom Python acquisition code.

### Surgery

Rats were anesthetized with isoflurane (5% induction; 1-3% maintenance) and standard stereotaxic procedures were used to implant 26 gauge dual cannula (1.5 center-to center distance; RWD, Shenzen, China) into the ventral tegmental area. Briefly, the scalp was cleaned with alternating alcohol and iodine scrubs and a midline incision was made. The skull was leveled between bregma and lambda and 4 skull screws were placed. The cannula was then implanted 1 mm above the intended infusion site in the ventral tegmental area (AP -5.6, ML ±0.75, DV -7.4) and secured to the skull and screws with dental cement. Rats received cefazolin (70 mg/kg, s.c) as a prophylactic antibiotic and carprofen (5 mg/kg, s.c.) to relieve pain. At all times other than during infusions flat cut dummy stylets were placed in the cannula. Rats recovered for 1 week following surgery with ad libitum access to food and water after which they returned to behavioral training and food restriction, if appropriate.

### Intracranial Infusions

One to two days before the first infusion, dummy stylets were removed and the patency of the cannula was confirmed by inserting an infuser into the guide cannula (31 gauge; RWD, Shenzen, China). Infusers extended 1 mm past the guide cannula. At test, rats were briefly anesthetized with isoflurane (5%) and received an intracranial infusion via 31 gauge infusers connected to 10 µL Hamilton syringes with PE50 tubing that were housed in a motorized pump (RWD, Shenzen, China). Rats received either saline, the nicotinic antagonist mecamylamine (300 mM), or the muscarinic antagonist scopolamine (350 mM). These doses of drugs were selected as they have been shown to elicit behavioral effects over 2-3 hours and were without locomotor impairment (Addy et al., 2015; Wickham et al., 2015; Nunes et al., 2019). Bilateral infusions of 0.3 µL of solution occurred over 1 minute, after which the infusers were left in place for 1 additional minute to allow for diffusion. After the infusion, dummies were replaced and rats were allowed to recover in their home cage. Anesthesia was used to prevent unmitigated stress from the infusion procedure and mitigate undue experimenter-induced variation in the experience for the rats across experiments (Narayanan et al., 2006; Erlich et al., 2015; Akhlaghpour et al., 2016; Palmer et al., 2024). Total anesthesia time was ∼4 minutes and rats recovered from anesthesia within 1 minute. All testing occurred 20 minutes after recovery. Order of infusions and testing order was counterbalanced across rats.

### Pavlovian Conditioning and Contextual Renewal

In daily 120 minute sessions hungry rats (n=22, 10 male, 12 female; food-restricted to 90% of free-feeding weight) learned to associate a 10 s auditory stimulus (CS+; white noise or 4.5 kHz tone) with the delivery of 0.1 mL of 10% sucrose at its offset. The other 10 s auditory stimulus not paired with sucrose delivery was assigned the CS- and was presented but never followed by reward. In each session there were 30 CS+ and 30 CS-trials in a pseudorandom order with an average 90 s variable intertrial interval. Rats were assigned to be trained in one of two conditioning chamber designs composed of unique flooring, lighting, and odor. One context was composed of strawberry odor, the stainless steel grid floor, and was illuminated by the houselight and the other context had an orange odor, a textured plastic floor, and was dark. The assignment of initial training context design was counterbalanced across rats and designated as Context A. After 14 training sessions, rats underwent 7 extinction sessions in the opposite chamber design, designated Context B. Task parameters were identical with the exception that no sucrose was delivered. After this rats were returned to Context A for a session with identical task parameters but no sucrose was delivered to assess contextual renewal. Rats then were retrained in Context A for 4 sessions, extinguished in Context B for 3 sessions, and tested for renewal in Context A again. We then retrained the rats in Context A and tested for the effect of acetylcholine manipulations in the ventral tegmental area to the performance of conditioned responding in Context A when reward was present. In this phase there were 1-2 days between tests. All rats consumed all reward delivered at the rewarded test. Rats in this experiment were assigned to pseudogroups and received 2-3 tests for renewal and during rewarded conditioning sessions in a within-subject design.

### Reward Consumption

Hungry rats (n=21; 9 male, 12 female; 90% food restriction) were allowed free access to two bottles in the modified homecage as described above. Rats were allowed 1 hour of access to 10% sucrose and water in ball-bearing sipper bottles. Rats were weighed before testing and bottles were weighed before and after each test to evaluate consumption. For a subset of rats, licking microstructure was assessed via a capacitive lickometer attached to the bottle containing 10% sucrose. After the test rats were returned to their homecage. At least 1 day without testing was between each test. The order of infusions were counterbalanced across rats. Rats were assigned to pseudogroups as above to limit the total number of infusions each rat received.

### Pavlovian-to-instrumental Transfer

Hungry rats (n=21; 12 male, 9 female; food-restricted to 85% of free-feeding body weight) were trained to first associate a 2-min auditory stimulus (CS+; white noise or 4.5 kHz tone) with the delivery of 0.1 mL of 20% sucrose on a random time 30s schedule (10-50 second range) during its presentation in 8 daily sessions as in (Collins et al., 2016). In each session there were 6 CS+ trials separated by a 4 minute average intertrial intertrial interval. After this rats were trained to press a lever for 0.1 mL of 20% sucrose. Initially rats received 1-3 sessions of training on a continuous reinforcement schedule until they earned 50 rewards in one hour. After this, rats transitioned to a variable interval schedule of reinforcement where reward was delivered at the first lever press following a variable delay. Rats received 1 session of a variable 15 s variable schedule, 1 session of a variable 30 s schedule, then 5 sessions of a variable 60 s schedule. These sessions were 30 minutes in length or ended if the rat earned 30 rewards. Rats then received surgery as described above.

After recovery rats had a session of Pavlovian retraining, and a session of habituation to the control cue (CS-) where the opposite 120 s auditory stimulus was presented 6 times on a 4 minute intertrial interval without reward. Rats then underwent lever-press retraining for 2 sessions and the day before testing had a 30 minute session of lever-press extinction to establish low press rates for test. In the tests of Pavlovian-to-instrumental transfer, rats were presented with the lever and pressing extinguished for 5 minutes after which the CS+ and CS-were presented 4 times each in a pseudorandom order with a fixed 4 minute intertrial interval. No reward was delivered at test. Between each test rats received 2 sessions of lever-press training, 1 session of Pavlovian conditioning, and 1 session of lever press extinction. Each rat was tested under each drug condition in a counterbalanced manner.

### Threat Conditioning

Before conditioning rats were randomly assigned to an infusion group (saline n=8, 4 male, 4 female; mecamylamine n=7, 3 male 4 female; scopolamine n=8, 4 male, 4 female). 20 minutes prior to conditioning rats received their respective infusion. Rats were trained in a distinct context (peppermint scented, wire grid floor, lights off, fan on, doors closed) to associate a 10 s 4.5 kHz tone with the delivery of a 2 s 1.0 mA foot shock through the grid floor at its offset. There was a 4 minute baseline period before the first cue-shock pairing afterwards there was a fixed 140 s intertrial interval. Rats remained in the chamber for 3 minutes after the last cue-shock pairing after which they returned to their homecage. The next day the rats returned to the chambers for a 10 minute test of context memory. The following day rats were returned to the chambers in a different design (lemon scented, plastic textured floor, lights on, fan off, doors open) and the tone CS+ was presented 15 times on a fixed 140 s intertrial interval after 3 a minute baseline period. Video recordings during all behavior sessions allowed for the offline scoring of freezing behavior. Freezing (Fanselow, 1980) was assessed by raters (LY & NSC) blind to group assignments during the baseline period as well as during the 10 s (2 s intervals) and a 60 s post shock period (6 s intervals) during conditioning and the cue test and over the 10 minute context test (6 s intervals) as adapted from (Hassell et al., 2025).

### Conditioned Suppression

Rats (n=22; 10 male; 12 female; 85% food restriction) were shaped to nosepoke for 0.1 mL of 20% sucrose on a variable interval 60 s schedule in 60 min sessions where sucrose delivery was signaled by the brief dimming of the houselight in the behavioral chamber for 0.5 s. Following nosepoke acquisition rats underwent 2 habituation sessions to the three 10 s auditory stimuli (white noise, 5 Hz clicker, 2.9 kHz tone) presented 4 times each in a random order with a variable 3 minute intertrial interval. The rats then were trained to associate one of the auditory stimuli with a 100% probability of a 0.5 s 0.3 mA footshock delivered 2 s after cue offset, one cue with a 25% probability of footshock, and the third cue was never associated with foot-shock. The identity of cue and probability pairings was counterbalanced across rats. There were 4 100% trials, 8 25% trials (2 shock, 6 no shock), and 4 0% trials in each session where each cue was presented randomly with a variable 3 minute intertrial interval. Task parameters were adapated from (Ray et al., 2020, 2022). Rats were trained for 12 sessions after which they received intracranial infusions in a series of 3 tests in a counterbalanced order with 1 day of retraining between each test. During each session fear was assessed by calculating a suppression index where the nosepoke rate during each cue was compared relative to a pre-cue baseline period of equivalent length over the total nose poke rate in the two periods. In this way a value of 1 would indicate complete suppression of nosepoking during the cue and -1 would be nosepoking solely during the cue.

### Progressive Ratio

Rats (n=18; 10 male; 8 female; 85% food restriction) were trained to nosepoke for 0.1 mL 20% sucrose on a variable interval 60s schedule in 60 minute sessions where sucrose delivery was signaled by the brief 0.5 s dimming of the houselight in the behavioral chamber. At test rats were shifted to a progressive ratio schedule as in (Richardson and Roberts, 1996) where each reward increased the subsequent amount of nosepokes required according to the formula:

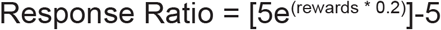

Response Ratio = [5e^(rewards * 0.2)^]-5

Which results in the response schedule: 1, 2, 4, 6, 9, 12, 15, 20, 25, 32, 40, 50, 62, 77, 95, 118, 145, 178, 219, 268, 328, 402, 492, 603, etc.

Rats were tested in a within subjects design and had 1 day off without testing between each progressive ratio test. The order of infusions was counterbalanced across rats. Sessions ended after 2 hours.

### Histology

Following completion of testing rats were deeply anesthetized with sodium pentobarbital and perfused with 4% paraformaldehyde. Brains were extracted and stored in 4% paraformaldehyde overnight then transferred to 30% sucrose in 0.1 M NaPB. 50 um thick coronal slices through the ventral tegmental area were collected on a cryostat and wet-mounted onto Fisher SuperFrost PLUS microscope slides. Sections were then coverslipped with DAPI mounting medium (Vectashield). Cannula positions were then mapped onto a standard rat brain atlas (Paxinos and Watson, 2007; Fig. S2). All numbers in the manuscript reflect final subject sizes following any potential exclusions for missed targeting.

### Statistical Analysis

Statistical analyses and graphs were produced in GraphPad Prism 11. For Pavlovian conditioning and Pavlovian-to-instrumental responding, responses (port entries, port time, or lever presses) during each cue was analyzed with respect to an equivalent pre-cue period or was normalized by subtracting responding during the pre-cue period from responses made during the cue. We made use of repeated measures one-way or two-way ANOVA when appropriate and for data from fear conditioning a between subjects ANOVA was conducted. Post-hoc and planned comparisons were performed with comparisons made between each drug condition and saline and not between each drug condition.

## ACKNOWLEDGEMENTS

The authors have no biomedical financial interest or potential conflicts of interest. We thank the members of the Fraser Lab for their comments throughout the development of this manuscript. This research was supported by funds from the Research Corporation for Science Advancement (Grant Number SA-MBC-2024-067a).

## Supplemental Figures

**Figure S1.**
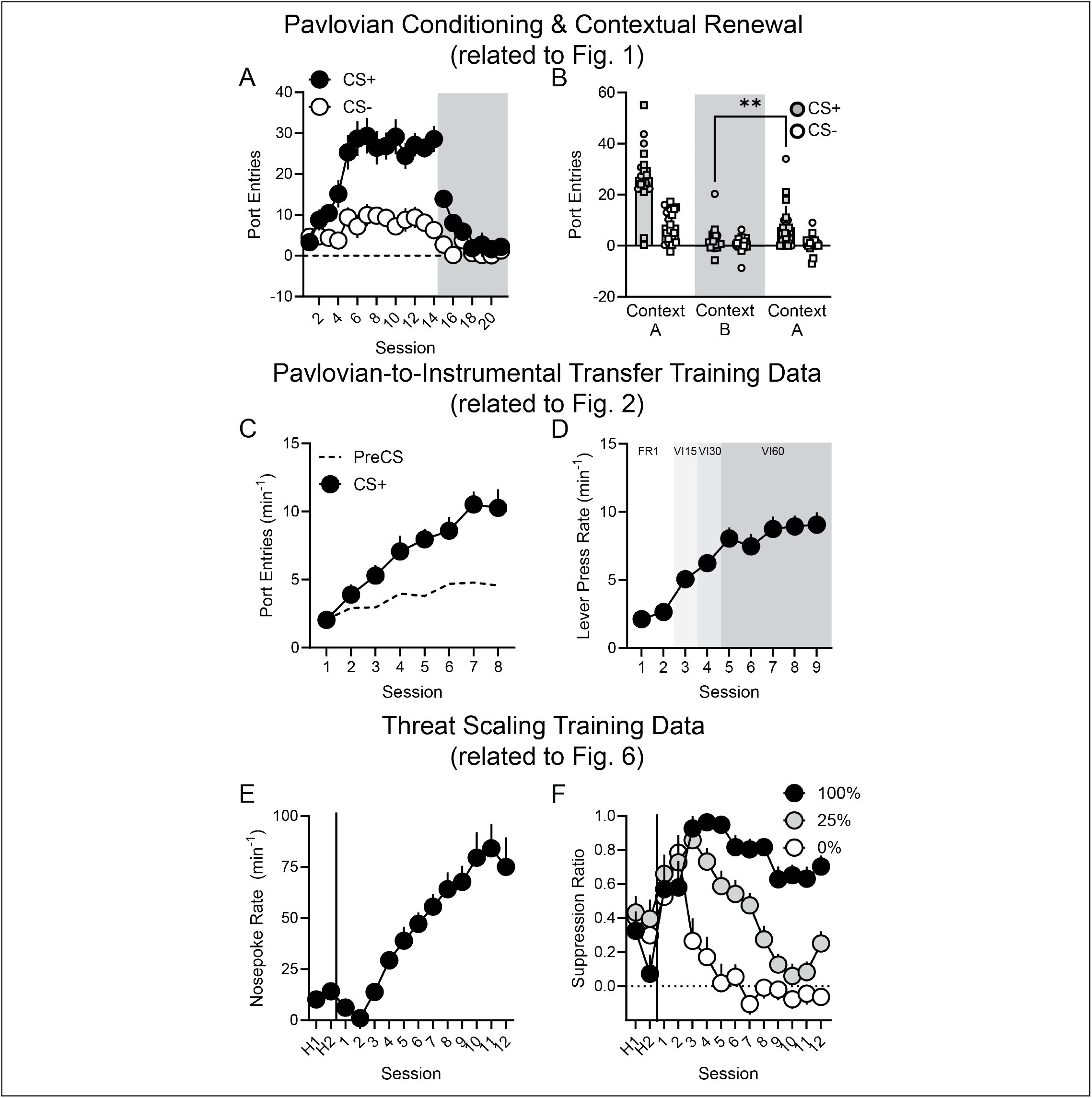
Training data for conditioning experiments. A) Rats acquired discriminated conditioned responding in Context A then were extinguished in Context B (shaded area denotes extinction). B) At test, all rats receiving saline demonstrated contextual renewal of reward-seeking as evidenced by an increased responding to the CS+ in Context A compared to in the extinction Context B (interaction F_(1,20)_=4.939, p=0.0379; post hoc p=0.0072). C) Responding in the period between cue onset and first reward delivery during the CS+ across training. D) Average lever press rates across training. E) Average nose poke response rate across training. F) Average suppression ratios across training. H1 and H2 indicate the two cue habituation sessions for threat scaling. Data presented represent mean + SEM. **p<0.01.

**Figure S2.**
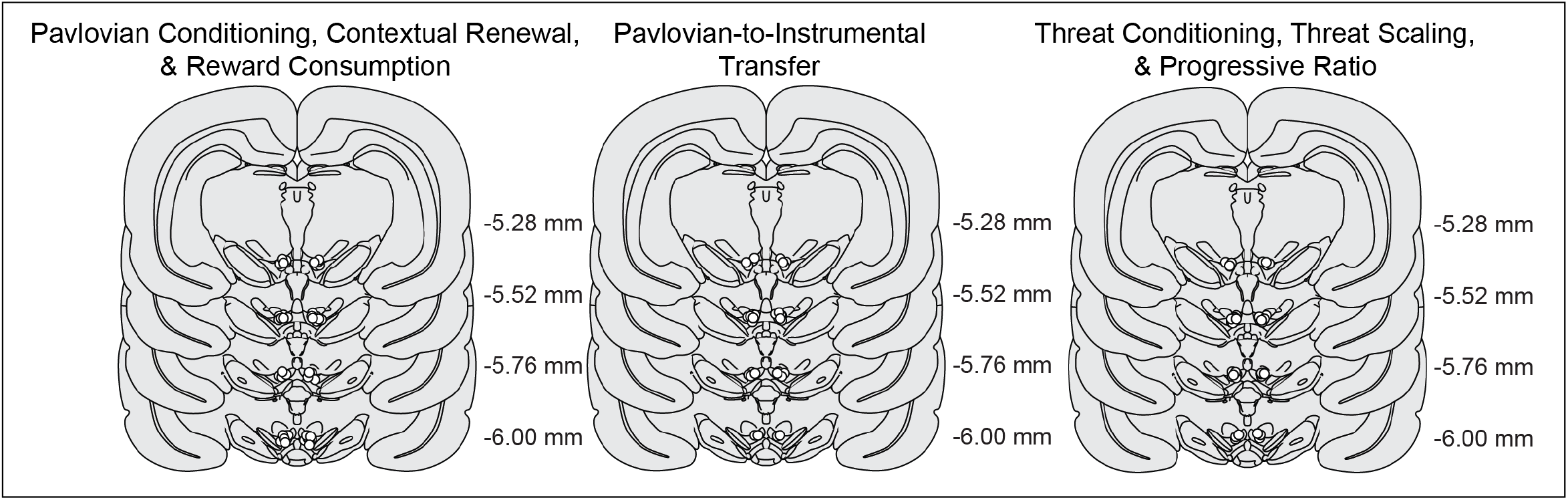
Cannula placements. Distribution of cannula placements for rats within each experiment mapped onto a standard brain atlas.

